# Biomechanical profiling of human epithelial corneal cells using atomic force microscopy

**DOI:** 10.1101/2025.08.07.669224

**Authors:** Taíse Tognon, Cleyton Alexandre Biffe, Carlos Alberto Rodrigues Costa, Priscila Cardoso Cristovam, Larissa Rigobeli da Rosa, Yara M. Michelacci, Eduardo Raad, Mauro Campos

## Abstract

Techniques for measuring the hardness of different substances have been described over the decades. In recent years, with the advancement of nanometric-scale evaluation technologies, interest in the biomechanical properties of cellular structures has grown. Atomic force microscopy (AFM) has emerged and evolved in this context, now featuring specialized modules for submerged specimens.

This study was conducted on live human corneal epithelial cells, using AFM indentation on specimens derived from primary cultures obtained from healthy post-mortem donor corneas via an eye bank. Hardness measurements were taken from different regions of the epithelial cells: central, paracentral, and peripheral. Differences in hardness were found in the peripheral regions compared to the central and paracentral zones, reviving the debate on epithelial architecture and function.

The measurement protocol presented is a pioneering approach in ophthalmology, and the findings differ from previously reported results for other cell types. Describing the biomechanical profile of ocular cells may greatly contribute to understanding the pathophysiology of ocular diseases, improving existing techniques, and serving as a foundation for new therapeutic strategies.

## Introduction

The human cornea is one of the key structures responsible for vision due to its role in light refraction. Its intricate architecture and anatomical features account for both its transparency and biological function. Several diseases can affect the corneal layers, altering their biomechanical properties. With advances in science and surgical techniques, new treatment modalities aimed at enhancing or restoring corneal hysteresis have emerged, notably corneal collagen crosslinking. However, there is still limited knowledge regarding the intrinsic stiffness of the cornea, which remains a significant limitation in refining current clinical procedures. This research was conceived to address this gap by investigating the stiffness of corneal cells.

Taking advantage of modern microscopy techniques—particularly atomic force microscopy (AFM)—it is now possible to conduct investigations at the nanometric scale and under in vivo-like conditions. Accordingly, this study aimed to establish a reliable protocol for assessing and characterizing the stiffness of live, normal human corneal epithelial cells using AFM. The present work is part of a broader investigation into the stiffness of both normal and pathological corneal cell types.

These assessments are grounded in the premise that corneas affected by structural deformities—such as keratoconus and pellucid marginal degeneration—are thinner and mechanically weaker than healthy human corneas. Nonetheless, to date, no studies have defined normal stiffness parameters for corneal cells.

It is well established that the biochemical alterations seen in such diseases can lead to structural changes that impact the cornea’s elastic modulus and overall mechanical stability. Characterizing the stiffness profile and mechanical behavior of normal corneal cells may contribute to the understanding of disease pathophysiology and inform the development or refinement of therapeutic strategies for corneal disorders.

## Materials and Methods

This innovative biomechanical study in ophthalmology was conducted through a collaboration between the LNNano laboratory, located at the Brazilian Center for Research in Energy and Materials (CNPEM) in Campinas, São Paulo, Brazil, and the Advanced Ocular Surface Center (CASO) of the Department of Ophthalmology and Visual Sciences at the Federal University of São Paulo – Paulista School of Medicine (UNIFESP/EPM) in São Paulo, Brazil.

### Acquisition of Cellular Specimens

Cell specimens were obtained from corneoscleral rims remaining from corneas used in transplantation, sourced from previously performed trephinations. Rims from 10 healthy corneas were used, provided by the Eye Bank of Hospital São Paulo (BOHSP). For tissue collection, written informed consent was obtained from the closest relatives of deceased donors at the time of organ donation, authorizing the use of the corneas for research purposes. These documents remain in the custody of the Eye Bank, complying with Brazilian regulatory standards, mainting privacy and total confidentialy. All procedures followed the principles established by the Declaration of Helsinki and strictly adhered to the regulations of the Eye Bank of Hospital São Paulo. The study was approved by the UNIFESP Research Ethics Committee (approval number 6.033.654) and by the Transplant Coordination Center of the State of São Paulo.

Inclusion criteria for the study were: corneas from donors aged 2 to 70 years, with a minimum endothelial cell count of 1800 cells/mm², and without corneal infiltrates or surgical scars. Exclusion criteria included opaque corneas, extensive epithelial defects, and diabetic donors. The corneoscleral buttons were prepared according to the standard operating protocols of the Eye Bank, preserved in Optisol-GS® (Bausch & Lomb, Rochester, NY, USA) or Eusol C® (Al.Chi.Mi.A. Srl, Viale Austria, Italy), maintained at 2 to 8°C, and used within 14 days of preservation. Morphological assessments and endothelial cell counts were performed during the preservation period to confirm tissue viability.

### *Ex-Vivo* Expansion of Corneal Epithelial Cells

The rims, obtained immediately after surgery, were placed in vials containing the aforementioned preservation media and transported to the CASO laboratory for cell culture. Under sterile conditions in a laminar flow hood, the corneoscleral rims were dissected and the endothelium scraped. The tissue fragments were then cut into pieces approximately 1–2 mm in size and seeded onto culture plates. These fragments, designated for epithelial cell culture, were placed in plates suitable for atomic force microscopy and incubated in a 1:1 mixture of Dulbecco’s Modified Eagle Medium and Ham’s F12 (Invitrogen, Grand Island, NY), supplemented with 10% fetal bovine serum, 2 ng/mL epidermal growth factor (Sigma-Aldrich, St. Louis, MO), 0.1 μg/mL cholera toxin (Sigma-Aldrich), 1 μg/mL recombinant human insulin (Sigma-Aldrich), 5 μg/mL hydrocortisone, penicillin-streptomycin, and amphotericin B (Sigma-Aldrich). Cells were incubated at 37°C in 5% carbon dioxide and 95% air, with medium changes every 2 to 3 days.

When initial outgrowth of cells from the explants was observed, the medium volume was increased to fully immerse the tissue fragments. No surface treatment or adhesion modification was performed on the plates to avoid artifacts in the atomic force measurements.

### Cell Transport and Experimental Preparation

Once the cultures reached 80–90% confluence, the culture medium was refreshed, and the cells were examined under an optical microscope. If viable and adherent cells were observed, the culture plates were transported from the CASO laboratory to LNNano in thermal boxes at ambient temperature, with minimal agitation. Upon arrival at LNNano, the plates were re-examined under an optical microscope and incubated at 37°C for at least 24 hours. After this period, under sterile conditions in a laminar flow hood, the DMEM/F12 culture medium was replaced with serum-free PBS solution. Viable cells exhibited a stellate and fusiform morphology on the plates. As they lost viability, they entered apoptosis, becoming rounded and bright, detaching rapidly from the surface. Before starting each experiment, and after each preparation step, the plates were evaluated under an optical microscope to confirm cell morphology and feasibility.

The above conditions ensured that cells on each plate remained viable and live for up to four hours—well beyond the average experiment duration of two hours.

### Atomic Force Microscopy

The structure of the atomic force microscope (AFM) consists of three essential components for measurements: a cantilever, a detection system for probe deflection, and a system for scanning and positioning the sample (see **Fig 1**).(1, 2)

**Figure 1.**
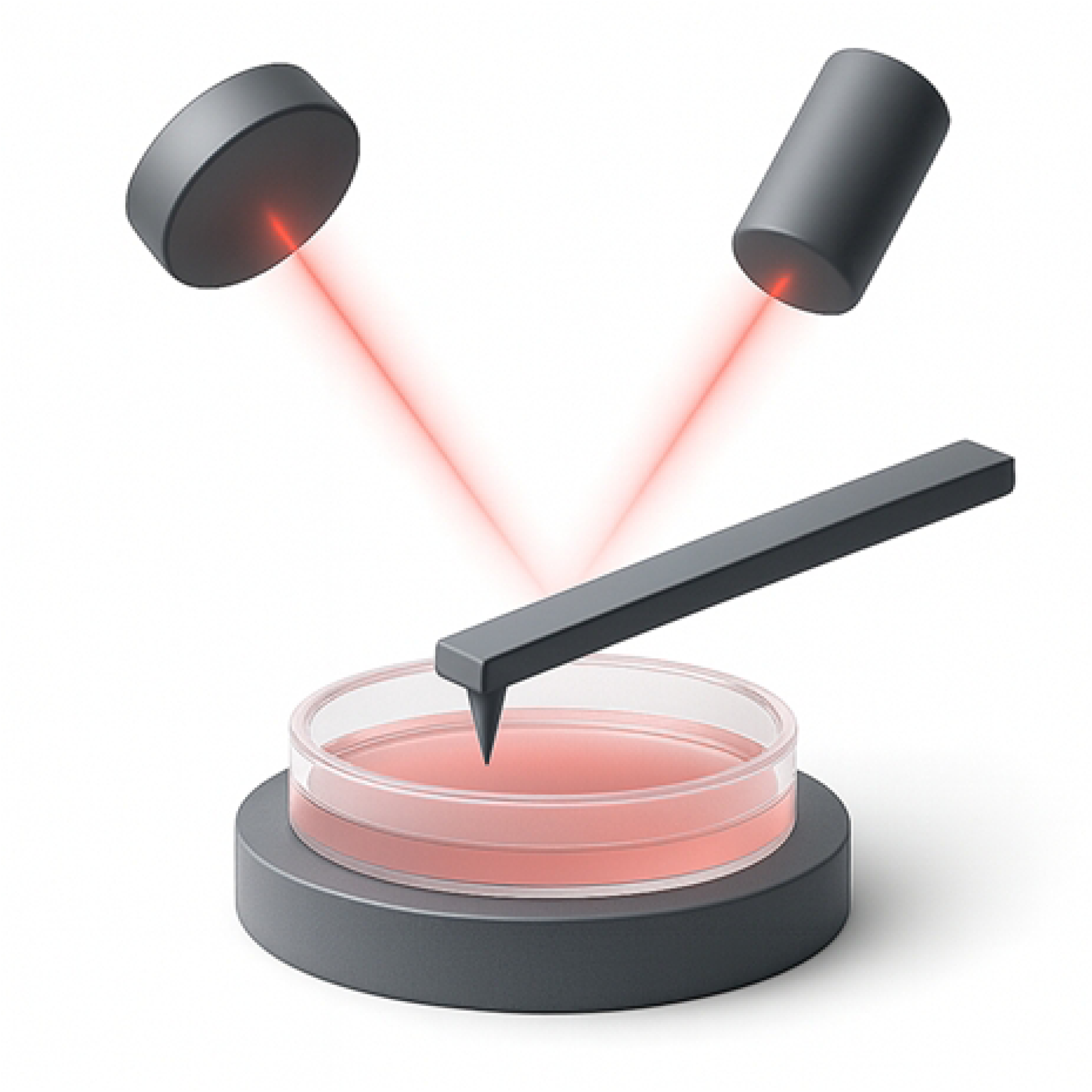
Schematic illustration of the atomic force microscope’s working principle, highlighting the LASER beam incident on the cantilever.

The operating principle of atomic force microscopy (AFM) turns possible the measurements with differents medium surrounding the cantilever (air, vacuum, or liquids).(2) The scanning and positioning system uses piezoelectric materials, enabling highly accurate positioning and scanning, and facilitating the handling and mounting of various samples.(2)

The cantilever is considered an important component of AFM.(2) It is lithographically fabricated from silicon or silicon nitride in the form of a long and flexible lever (rectangular or triangular), with a probe attached to its free end.(2) The mechanical properties of a cantilever are characterized by its corresponding spring constants, which for live cells typically range from 0.01 to 0.1 newtons per meter (N/m).(2)

As for the AFM probe tips, they may have various shapes: pyramidal, conical, or spherical.(1,2) For elastic mode measurements in live cells, the most suitable tips are those with an spherical shape and micrometric diameter, as they do not induce excessive pressure at the contact area—thus minimizing the risk of cell rupture from sharp tips or imprecise measurements—and they more appropriately satisfy the assumptions of the Hertz model, which is commonly used for estimating the Young’s modulus.(2)

In AFM, the Young’s modulus is a calculus of elasticity of a material based on the stiffness measure, indicating how it resists to elastic deformation when subjected to a force, and can be measured by analyzing the deformation of the sample surface under an applied force with the cantilever tip. The Young’s modulus measured by AFM is widely used in the study of cells, biological tissues, polymer, and materials characterization.

Measurements of elasticity are obtained through the interaction between the probe and the sample (in this case, the cell), which causes a deflection in the cantilever.(2) The most used method for detecting this deflection movement employs an optical system composed of a LASER and a photodetector.(2)

The atomic force microscope used in this study was the Bruker BioAFM model (NanoWizard4) (JPK/Bruker Corporation, Berlim, Germany) (see Box 1 for technical specifications). The equipment is in a dedicated room, fully equipped for such analyses, with continuous control of temperature, humidity and isolation for mechanical and acoustic vibration.

##### Box 1. Technical specifications of the atomic force microscope used in this study

###### Specifications of the Bruker BioAFM Atomic Force Microscope (NanoWizard4 and CellHesion200 models)

- NanoWizard XY drive range: 100 µm × 100 µm;
- NanoWizard Z drive range: 15 µm;
- CellHesion200 Z drive range: 100 µm;
- Motorized XY sample stage: 20 mm × 20 mm;
- Cytosurge pressure control system (from -1 bar to 5 bar);
- Temperature controller for Petri dish (ambient to 60°C);
- Zeiss Observer 5 inverted optical microscope;
- Objective lenses with magnifications of 10×, 20×, 40×, and 63×;
- Fluorescence system with DAPI, GFP, TX, and DsRed filters;
- Phase contrast and DIC filters;
- Top-view optical microscope with variable objective lens, up to 7× magnification.

In this study, after the culture plate was installed on the AFM system, wide-area scans ("rasters") were performed for surface mapping, cell localization, and morphological analysis through direct visualization using the integrated optical microscope. For measurements to be valid, the cells need to be non-confluent—that is, isolated, alive, and viable.

Initially, fixed cells were measured in liquid to analisys the height profile of this type of cell. For these topography images, the Quantitative Image (QI) mode was used, with the probe PFQNM-LC-A-CAL (Bruker), a pre calibrated cantilever (nominal spring constante of 0.1N/m) with a parabolic tip shape with 70nm radius. The images were acquired with 256x256 pixels of resolution and the scanning area depends on the cell shape.

Each cell was segmented into regions (based on what was visible on the microscope screen): two peripheral regions (upper and lower), two paracentral regions (upper and lower), and one central region. Five measurements were obtained from each of these regions, and each measurement was repeated three times at the same point to ensure internal control and validation. **Fig 2** illustrates photographic documentation from the optical microscope screen attached to the atomic force microscope during the acquisition process.

**Figure 2.**
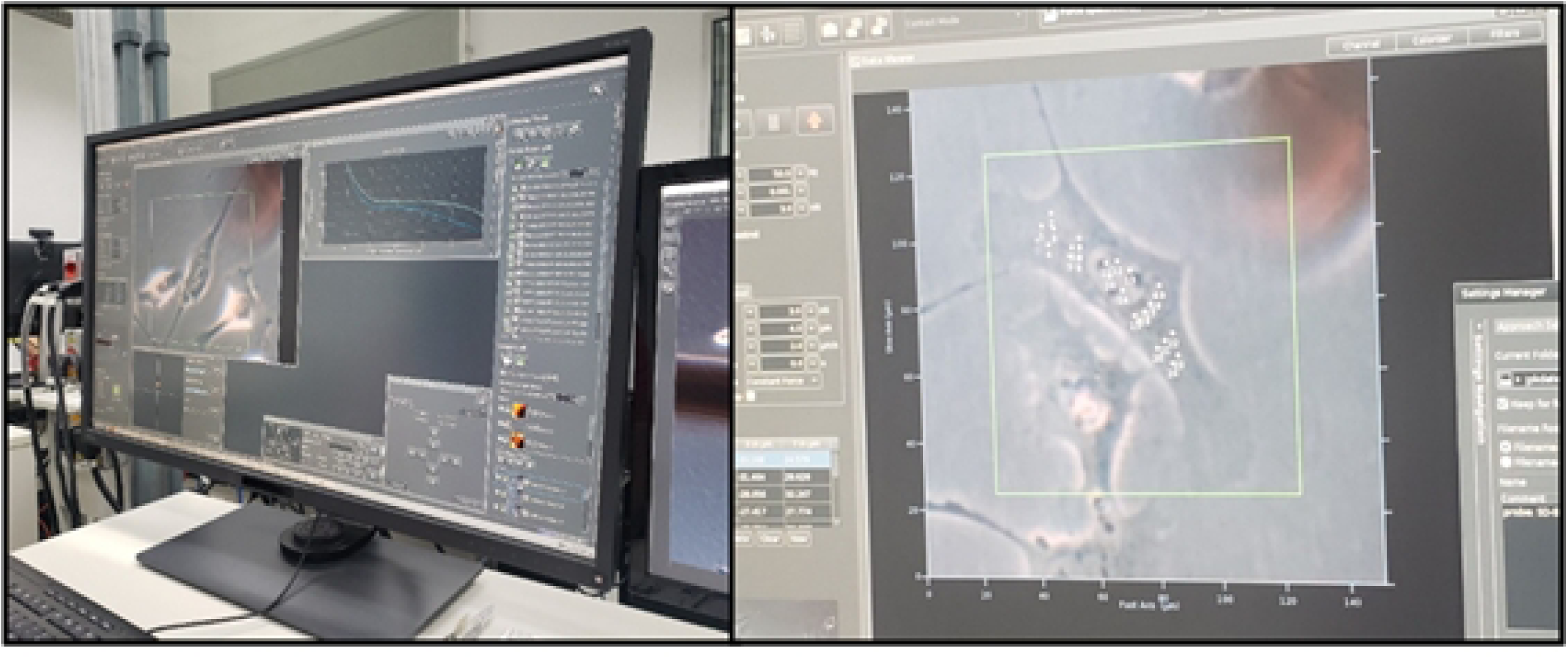
Photographic documentation of the normal corneal epithelial cell as visualized through the optical microscope coupled to the atomic force microscope, and the distribution of measurements according to cellular regions.

The atomic force microscopy acquisition parameters included force–distance curves with a Z length of 3 micrometers and a maximum indentation force of 3 nanonewtons, Z spped of 2 micrometers per second. Prior to the start of the experiments, the AFM probe model SD-SPHERE-Cont-L (Nanosensors Corporation, Switzerland), with a spherical tip (2 µm radius) and a nominal spring constant of 0.2 N/m, was carefully selected under optical microscopy guidance, aligned and calibrated accordingly, indenting the probe against a harder surface than the probe’s cantilever (saphire sample), and all measurements were performed in force spectroscope mode. For the assessment of cellular elasticity, the spherical Hertz model was adopted to estimate Young’s modulus.

The microscope was calibrated in liquid medium and remained powered on continuously for one week using the same probe. When not in active use, the probe was kept immersed in a dish with deionized water, under the same environmental conditions—temperature and humidity control— as the laboratory room, to maintain consistent parameters.

No substance for adhesion of epithelial cells to the plates was used, in order not to interfere with the measurements, and the acquisitions were all performed on live cells.

All stiffness measurements were recorded as force-distance curves, which were then converted into numerical values with JPK Data Processing software (JPK/Bruker). Cell topographies were also generated in graphic format and converted into visual images by the proprietary AFM software for optical and AFM overview, and by the open-source software Gwyddion for 3D view. Experimental data were saved within the microscope software and additionally backed up by the LNNano laboratory in a separate database, later converted into Excel spreadsheets and sent to the principal investigator.

## Statistical Methodology

Initially, the data were analyzed descriptively using summary measures (mean, quartiles, minimum, maximum, and standard deviation).

The reproducibility of the three replicated stiffness measurements for each region and position was assessed using intraclass correlation. Intraclass correlation quantifies overall agreement at the individual level across repeated measurements. Values below 0.5, between 0.5 and 0.75, between 0.75 and 0.9, and above 0.9 indicate poor, moderate, good, and excellent reproducibility, respectively.(3)

Mean stiffness values by region and position were compared using the Kruskal–Wallis test due to the non-normal distribution of stiffness, as verified using the Kolmogorov–Smirnov test. When significant differences were detected by the Kruskal–Wallis test, distinct groups of means were identified through Dunn–Bonferroni multiple comparisons to preserve the global significance level. In this comparison, the mean of the three replicates was used to calculate the regional and positional stiffness.

For all statistical tests, a significance level of 5% was adopted. Analyses were conducted using the statistical software package SPSS 20.0 (IBM Corp. Released 2011. IBM SPSS Statistics for Windows, Version 20.0. Armonk, NY: IBM Corp.).

## Ethics Statement

This study was approved by the Research Ethics Committee of the Federal University of São Paulo (CAAE: 04604418.1.0000.5505), as well as LNNano Laboratory, CASO Laboratory, UNIFESP Department of Ophthalmology and Visual Sciences, Eye Bank of Hospital São Paulo and Transplant Coordination Center of the State of São Paulo. All experimental procedures adhere to the Declaration of Helsinki and institutional guidelines for biomedical research involving human tissue.

## Results

Initially, some cells were randomly selected and topography measurements performed showed that the average maximum height and standard deviation of these cells is 6,9±1,3μm, as shown in an example in **Fig 3**.

**Figure 3.**
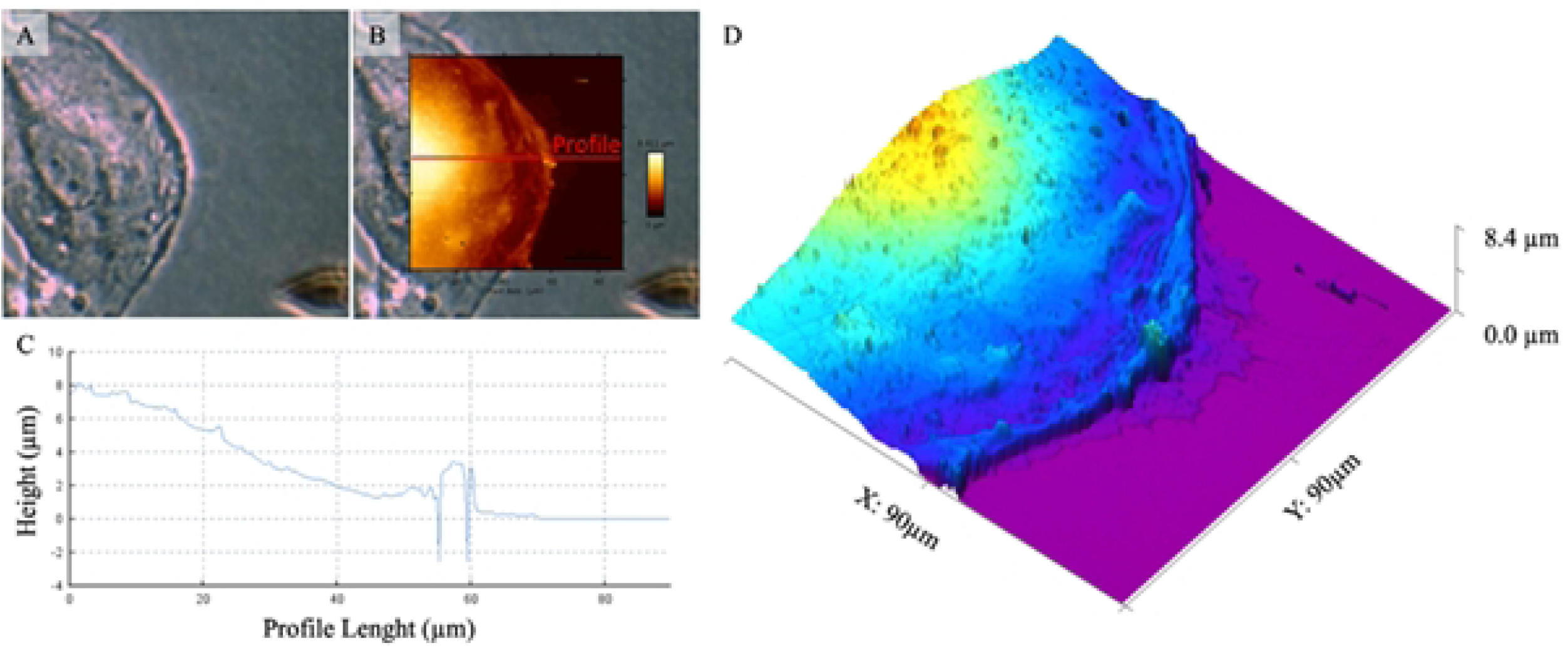
Optical and topographic image of an epithelial corneal cell.

Optical phase contrast image (A) can be overviewed with the AFM image (B), where it indicates (red line) an extracted height profile (C). The topography details of the cell can be observed in the 3D view (D), which false color gradient also represents the height information and helps to give contrast to the surface details.

A total of 60 cells were randomly selected for stiffness measurement, obtained from 10 corneas. Three measurements were performed at each point, with five points assessed across five different macroscopic regions of each cell, totaling 900 individual measurements.

According to **Table 1**, excellent reproducibility was observed for all regions: central (ICC = 0.941; 95% CI: 0.928 to 0.952; p < 0.001), inferior paracentral (ICC = 0.943; 95% CI: 0.931 to 0.954; p < 0.001), superior paracentral (ICC = 0.920; 95% CI: 0.903 to 0.935; p < 0.001), inferior peripheral (ICC = 0.914; 95% CI: 0.895 to 0.930; p < 0.001), and superior peripheral (ICC = 0.951; 95% CI: 0.940 to 0.960; p < 0.001). Regarding positional reproducibility, excellent or very good levels were found — intraclass correlation coefficients ranged from 0.807 to 0.985.

**Table 1.** Intraclass correlation for stiffness by region and position.

|  | Intraclass Correlation Coefficient (ICC) | 95% CI | N | p |
| --- | --- | --- | --- | --- |
| **Central Region** | **0,941** | **0,928 a 0,952** | **293** | **<0,001** |
| Position 1 | 0,870 | 0,798 a 0,920 | 56 | <0,001 |
| Position 2 | 0,955 | 0,930 a 0,971 | 59 | <0,001 |
| Position 3 | 0,971 | 0,956 a 0,982 | 60 | <0,001 |
| Position 4 | 0,946 | 0,917 a 0,966 | 59 | <0,001 |
| Position 5 | 0,955 | 0,931 a 0,972 | 59 | <0,001 |
| **Inferior Paracentral Region** | **0,943** | **0,931 a 0,954** | **297** | **<0,001** |
| Position 1 | 0,976 | 0,962 a 0,985 | 60 | <0,001 |
| Position 2 | 0,922 | 0,880 a 0,951 | 59 | <0,001 |
| Position 3 | 0,946 | 0,917 a 0,966 | 59 | <0,001 |
| Position 4 | 0,946 | 0,917 a 0,966 | 59 | <0,001 |
| Position 5 | 0,932 | 0,896 a 0,957 | 60 | <0,001 |
| **Superior Paracentral Region** | **0,920** | **0,903 a 0,935** | **290** | **<0,001** |
| Position 1 | 0,964 | 0,944 a 0,978 | 56 | <0,001 |
| Position 2 | 0,979 | 0,968 a 0,987 | 59 | <0,001 |
| Position 3 | 0,807 | 0,703 a 0,879 | 59 | <0,001 |
| Position 4 | 0,887 | 0,826 a 0,929 | 58 | <0,001 |
| Position 5 | 0,985 | 0,976 a 0,990 | 58 | <0,001 |
| **Inferior Peripheral Region** | **0,914** | **0,895 a 0,930** | **292** | **<0,001** |
| Position 1 | 0,983 | 0,974 a 0,989 | 60 | <0,001 |
| Position 2 | 0,982 | 0,972 a 0,989 | 57 | 0,048 |
| Position 3 | 0,888 | 0,826 a 0,930 | 57 | <0,001 |
| Position 4 | 0,960 | 0,939 a 0,975 | 59 | <0,001 |
| Position 5 | 0,973 | 0,958 a 0,983 | 59 | <0,001 |
| **Superior Peripheral Region** | **0,951** | **0,940 a 0,960** | **298** | **<0,001** |
| Position 1 | 0,905 | 0,855 a 0,940 | 60 | <0,001 |
| Position 2 | 0,964 | 0,946 a 0,978 | 60 | <0,001 |
| Position 3 | 0,942 | 0,911 a 0,963 | 60 | <0,001 |
| Position 4 | 0,976 | 0,963 a 0,985 | 60 | <0,001 |
| Position 5 | 0,979 | 0,968 a 0,987 | 58 | <0,001 |

95% CI=95% Confidence Interval.
N=Number of tests.
p=p-value.

In the following analyses shown in **Table 2**, stiffness by position and region was evaluated using the means of the three replicate stiffness measurements. As shown, no significant differences in mean stiffness were observed between positions across all regions.

**Table 2.**
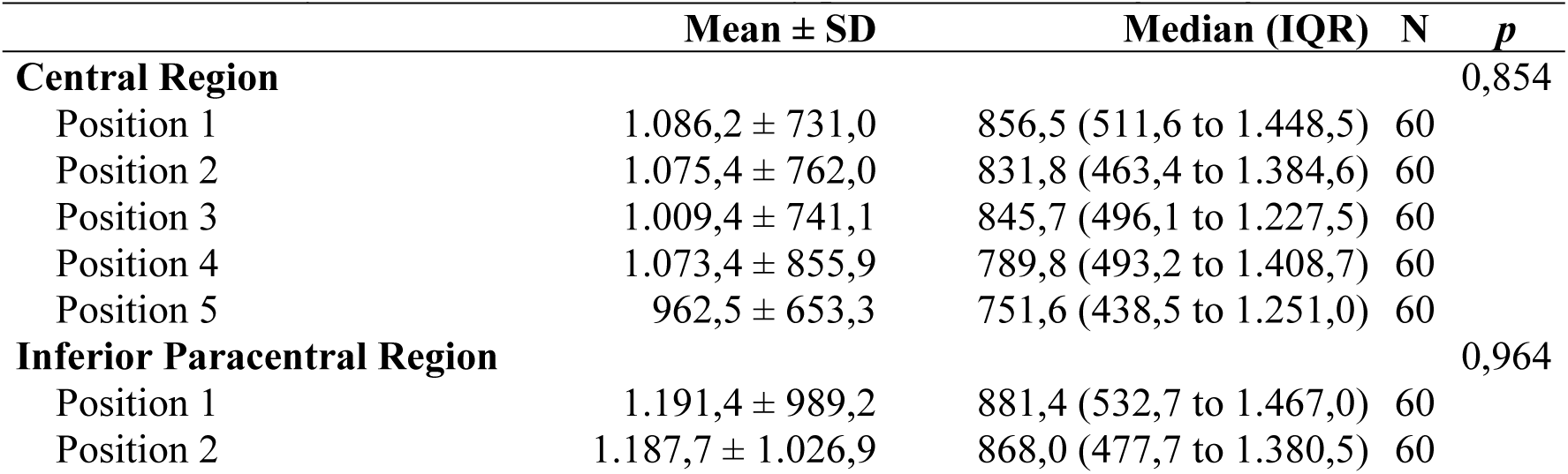
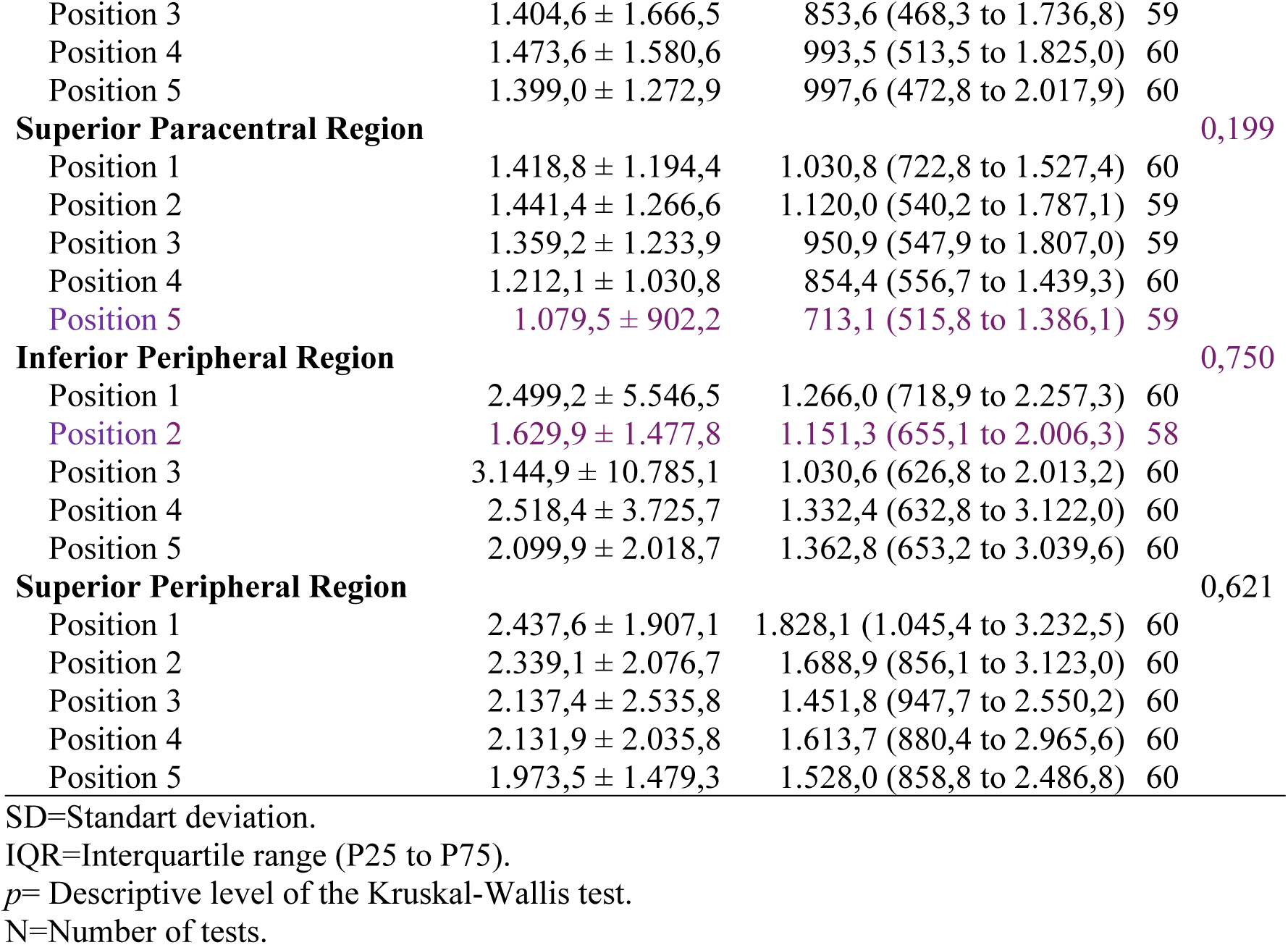
Summary measures of cell stiffness by position, according to region.

In **Table 3**, significant differences in mean stiffness across regions were observed (p < 0.001). Specifically, the mean stiffness values of the inferior and superior peripheral regions were similar to each other and higher than those of the other regions, which were also similar among themselves.

**Table 3.** Summary measures of stiffness by region.

| Region | Mean ± SD | Median (IQR) | N | p |
| --- | --- | --- | --- | --- |
| Central<sup>‡</sup> | 1.041,4 ± 748,0 | 802,1 (488,3 to 1.368,9) | 300 | <0,001 |
| Inferior Paracentral<sup>‡</sup> | 1.331,0 ± 1.331,4 | 909,1 (497,9 to 1.681,5) | 299 |  |
| Superior Paracentral<sup>‡</sup> | 1.302,3 ± 1.134,5 | 952,7 (565,6 to 1.549,2) | 297 |  |
| Inferior Peripheral<sup>§</sup> | 2.383,5 ± 5.784,0 | 1.189,9 (656,7 to 2.623,0) | 298 |  |
| Superior Peripheral<sup>§</sup> | 2.203,9 ± 2.028,4 | 1.649,1 (909,0 to 2.879,4) | 300 |  |

SD=Standart deviation.
IQR=Interquartile range (P25 to P75).
p= Descriptive level of the Kruskal-Wallis test.
N=Number of tests.
‡ and § indicate statistically different means according to Dunn-Bonferroni post hoc comparisons.

## Discussion

With recent advances in nanotechnology, several innovative experimental techniques have been developed to mechanically probe individual cells using applied forces and displacements.(4) Atomic force microscopy (AFM) stands out in this regard, as it is a surface-scanning analytical technique that enables the detection and measurement of a sample’s topographic features.(5–7) It proves to be an excellent tool for cellular biology applications since it produces nanoscale results and can be performed on a variety of samples, including those in liquid environments—allowing for detailed analysis of cells in their natural state, for example.(5,7–12)

The basic idea of AFM is to use mechanical scanning with probes over a sample surface, through the interaction between the probe tip and the sample itself.(11) Compared to conventional optical microscopy, AFM provides better spatial resolution (down to sub-nanometer levels), which allows mapping of the distribution of even isolated molecules.(11,13)

Electron scanning and transmission electron microscopy also offer high resolution, but the complexity of sample preparation (e.g., chemical fixation, dehydration, metal coating, and ultrathin sectioning) can substantially distort the specimens.(11) Since the AFM probe directly contacts the sample, minimal or even no preparation is required in some cases.(11) Moreover, AFM is one of the only techniques capable of providing direct structural, mechanical, and functional information at high resolution.(12)

For these reasons, the research team opted to use AFM to measure the stiffness of cells— enabling the acquisition of ‘real’, in vivo measurements. In this study, we present the use of this tool on corneal epithelial cells derived from primary cultures, representing an innovation in the field of ophthalmology.

Although innovative in ophthalmology, AFM analysis of living cells has already been applied in other medical fields, such as the study of prokaryotic cells (bacteria, fungi) and eukaryotic cells (tumor cells, myocardial, skeletal, and endothelial cells).(1,10,15–19) The emerging use of AFM has made it a valuable tool not only for topographic surface analysis under physiological conditions but also for localized measurement of mechanical properties of samples.(15)

Regarding the study of cell stiffness and elasticity, AFM techniques operate in one dimension, usually coupled with two-dimensional imaging.(5,15,16) For assessing these mechanical properties, the probe indents the sample at a predetermined set point, generating a force response proportional to the cantilever deflection and the distance traveled by the probe tip.(17) These force-distance curves reflect how the sample responds under load and can be used to calculate the Young’s modulus (a quantitative measure of elasticity) and other mechanical properties.(17)

Mechanical stress and resulting deformation play a central role in cellular contraction, spreading, crawling, invasion, wound healing, and cell division, and have been implicated in protein regulation, DNA synthesis, and programmed cell death.(20) These properties are largely due to the cytoskeleton and its dynamic behavior, which functions as a sol-gel transition system, allowing it to behave as a fluid in some situations (sol phase) and as a solid in others (gel phase).(20)

To examine and describe the mechanical properties of cells, a variety of techniques exist that consider different forces and timescales, including magnetic twisting cytometry, micropipette aspiration, particle tracking, optical tweezers, microplate rheology, and AFM, as previously mentioned.(21,22)

Among these, AFM force-indentation experiments are frequently performed to access a wide force range and to generate more accurate maps of cellular elasticity and adhesion properties.(22) However, the elastic response of confluent cells to indentation is commonly modeled through continuous mechanical contact frameworks.(22)

When using this model, many features and characteristics of the sample may be overlooked in favor of obtaining a single parameter: the Young’s modulus of the material under study.(22) Although the Young’s modulus allows for quick comparisons, it does not capture the complex architecture and active contractility of live cells, and thus only provides a qualitative representation of cellular stiffness.(19,22) This was the modeling approach adopted in our study—evaluating isolated cells in liquid medium via indentation using Young’s modulus—and therein lies one of our limitations.

Epithelial cells do not naturally exist in isolation but form dense, continuous sheets at interfaces, with highly dynamic cell–cell junctions.(22) These cell layers are often exposed to various mechanical challenges due to osmotic pressure changes, chemical stimuli, or external stress.(22) Describing biomechanical properties in live isolated cells, as done in our experiment, may provide initial insights and help guide future investigations, but studies must progress toward tissue-level interface analysis.

Eukaryotic cells are composed of the plasma membrane, cytoplasm, and a nucleus.(4) The cytoplasm can be further divided into a dense outer ectoplasm, primarily responsible for cell movement, and a granule-free, less dense endoplasm, which contains most cellular structures.(4) Thus, the interior of a cell is far from homogeneous in reality.(4)

Furthermore, the nucleus has been shown to be significantly stiffer and more viscous than the surrounding cytoplasm.(4,23–25) Studies using homogeneous liquid models, including Newtonian and Maxwellian models, have revealed that apparent viscosity and stiffness continuously vary with the degree of deformation.(4,23,26,27) Therefore, a more complex model, comprising heterogeneous components, is necessary to accurately reflect the physical composition of the cell.(4)

In our study using the AFM indentation model, no differences in stiffness were observed between central and paracentral regions of the cells—contrary to findings from earlier studies using other techniques and different cell types.

The mechanics of the plasma membrane and cytoskeleton play a fundamental role in the cellular response to mechanical stimuli.(22) While the mechanics of the membrane dominate under high tension due to the inextensibility of the lipid bilayer, the actomyosin cytoskeleton is responsible for responses to small deformations and active stress components.(22)

The cytoskeleton is an active gel with transient crosslinking that supports passive stress but also generates tension via motor activity, consuming chemical energy, and through actin filament polymerization and depolymerization.(22) Unlike synthetic polymer networks, the cellular cytoskeleton is a highly dynamic system due to the binding/unbinding kinetics of crosslinking proteins and the polymerization cycles of the filaments themselves.(21)

One of the most striking features of the cytoskeleton is its ability to perform antagonistic mechanical functions—supporting large elastic stresses while also retaining the ability to flow, such as during cell migration.(21) Notably, this biopolymer gel can generate internal tension by consuming chemical energy; this system operates far from equilibrium, making the study of cytoskeletal mechanics both fascinating and challenging.(21)

Most of the mechanical properties of epithelial cells originate from a specific region of the cytoskeleton—a thin actin-myosin mesh layer known as the cell cortex(22), located adjacent to the cell membrane (as shown in **Fig 4**). This cortex forms a dense active gel with a mesh size of a few dozen nanometers, consisting of transiently crosslinked actin filaments.(22) The tension generated by this active cortex produces a pressure difference responsible for bleb formation and pseudopod extension, prerequisites for cell locomotion.(22)

**Figure 4.**
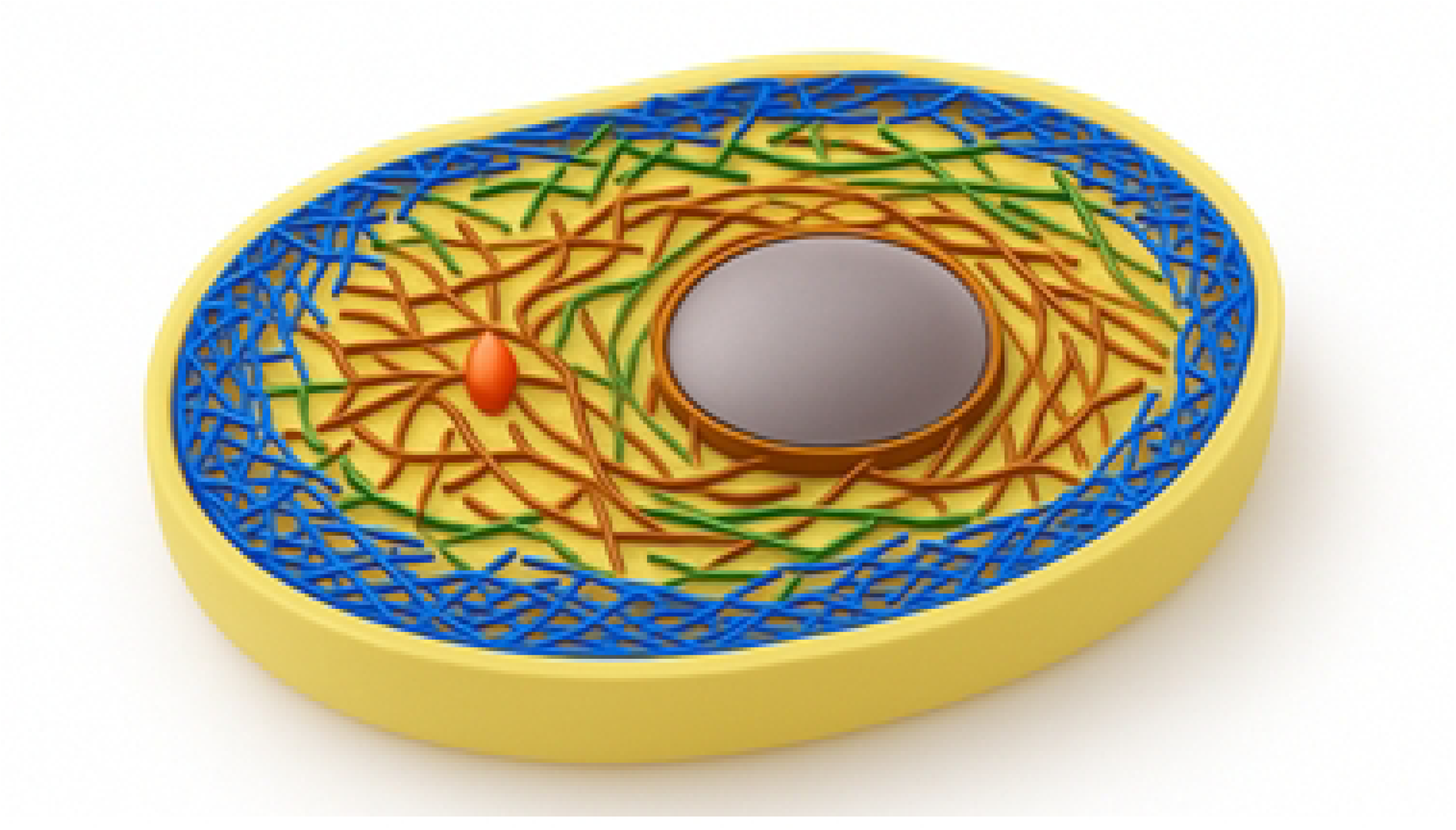
Schematic representation of the organization of cytoskeletal filaments in a eukaryotic cell.

Blue filaments depict the cortical actin network, the orange structure marks the centrosome, brown filaments correspond to intermediate filaments, and green filaments represent microtubules. The nucleus is paracentrally positioned.

In addition to cortical tension being the primary source of mechanical force in suspended cells, plasma membrane tension is recognized as an important regulator of many cellular processes that involve changes in membrane surface area, including cell adhesion, migration, mitosis, endocytosis, exocytosis, membrane repair, osmoregulation, and cell spreading.(22,28)

Thus, we believe that the differences in stiffness observed in various regions of normal corneal epithelial cells in our study are attributable to this intricate cortical actin mesh. This actomyosin network is contractile and generates cytoskeletal prestress, which enables the cell to stiffen through the production of intracellular forces.(22)

In the peripheral areas, the concentration and activity of this component tend to be higher than in the central and paracentral regions. This is because peripheral regions of cells are where pseudopod formation and other previously described activities primarily occur. In contrast, the central and paracentral regions of the cell are predominantly composed of intermediate filaments and microtubules. The network here is less intricate, has reduced mechanical strength, and contains a higher density of organelles and the nucleus.

Moreover, it has been proposed that the elastic response of cells is a function of environmental stimuli.(4,18,22,29) For example, epithelial cells cultured on porous or soft substrates appear softer than those grown on flat or rigid substrates.(22,29)

Living human cells are continuously subjected to mechanical stimuli throughout life, originating both from the external environment and from internal physiological conditions.(4)

Depending on the magnitude, direction, and distribution of these mechanical stimuli, cells may respond in different ways: shear stress on endothelial cells triggers hormone release, intracellular calcium signaling, and stiffening of the cells through cytoskeletal rearrangement; compression may influence the cell’s shape and function as well.(4)

Additionally, tensile stretching of the cell’s substrate can alter both motility and orientation of the cell.(4) Therefore, understanding how a cell mechanically responds to physical loads is a critical first step in further investigating how mechanical signals are eventually converted into biological and chemical responses within the cell.(4)

This underlines the importance of our study, which is pioneering in its description of cells of ocular origin. Understanding the properties of these isolated cells, establishing normal parameters, and subsequently investigating pathological behaviors will help improve existing treatments (such as corneal collagen crosslinking) and aid in developing new therapies for corneal biomechanical diseases.

In addition to mechanical loads generated inside or outside the body, many chemical substances are also known to increase or decrease the mechanical properties of living cells.(4) For instance, chemotactic agents can increase neutrophil stiffness; cytochalasin D and latrunculin B can disrupt actin filaments in the cytoskeleton and negatively impact cellular stiffness; colchicine can disrupt microtubules in neutrophils’ cytoskeleton.(4)

Therefore, the mechanical properties of certain cell types may potentially be used to quantitatively reflect the state of their structural and cytoskeletal health. This could indeed be useful for possible applications in clinical diagnostics and even therapy for certain types of diseases.

Finally, it is known that the mechanical properties of individual cells can determine the structural integrity of entire tissues, resulting from mechanical interactions between the cells and the surrounding extracellular matrix.(4) Conversely, mechanical loads applied at the tissue level are transmitted to individual cells and can influence their physiological functions.(4)

Although some progress has been made in developing tissue-level mechanics, studying the mechanics of individual cells remains a challenge—particularly when considering the living and dynamic nature of the cell.(4) Thus, our research has the limitations of being an experiment involving cells cultured in an environment that differs from their natural tissue context and of employing the Hertz model for cell stiffness verification.

Future perspectives in this field will involve studies on various types of isolated, living human cells, within intact tissues, and comparisons between normal and pathological specimens.

## Authors’ Contributions

**Taíse Tognon**: Conceptualization, Methodology, Data Curation, Formal Analysis, Investigation, Writing – Original Draft.

**Cleyton Alexandre Biffe:** Conceptualization, Methodology, Formal Analysis, Investigation and Revision.

**Carlos Alberto Rodrigues Costa:** Methodology, Formal Analysis, Investigation, and Supervision.

**Priscila Cardoso Cristovam:** Methodology, Formal Analysis, Investigation, and Revision.

**Larissa Rigobeli da Rosa:** Methodology, Investigation and Revision.

**Yara M. Michelacci:** Conceptualization and Revision.

**Eduardo Raad:** Conceptualization.

**Mauro Campos:** Conceptualization, Methodology, Formal Analysis, Investigation, Revision and Supervision.

## Data Availability

All relevant data is included in the manuscript. Additional raw datasets are available from the corresponding author upon reasonable request.

## Ethics Statement

This study was approved by the Research Ethics Committee of the Federal University of São Paulo (CAAE: 04604418.1.0000.5505). All experimental procedures adhere to the Declaration of Helsinki and institutional guidelines for biomedical research involving human tissue.

## Acknowledgments

The authors thank the staff of the Department of Ophthalmology and Visual Science at the Federal University of São Paulo for their technical support and collaboration.

